# Catalytic regulation of CoA transferase by an NAD^+^-sensing accessory protein and protein acetylation

**DOI:** 10.1101/2025.10.31.685773

**Authors:** Ayako Yoshida, Hiroyuki Yamamoto, Tomoko Miyata, Takeo Tomita, Minoru Yoshida, Keiichi Namba, Saori Kosono, Makoto Nishiyama

**Affiliations:** Graduate School of Agricultural and Life Sciences, The University of Tokyo, 1-1-1 Yayoi, Bunkyo-ku, Tokyo 113-8657, Japan; Collaborative Research Institute for Innovative Microbiology, The University of Tokyo, Tokyo, 1-1-1 Yayoi, Bunkyo-ku, Tokyo 113-8657, Japan; Graduate School of Frontier Biosciences, Osaka University, Suita, Osaka 565-0871, Japan; JEOL YOKOGUSHI Research Alliance Laboratories, Osaka University, Suita, Osaka 565-0871, Japan; RIKEN Center for Sustainable Resource Science, 2-1 Hirosawa, Wako, Saitama 351-0198, Japan

**Keywords:** alanine dehydrogenase, CoA transferase, NAD^+^/NADH ratio, protein acetylation, pseudoenzyme

## Abstract

CoA transferases play essential roles in short-chain fatty acid metabolism by catalyzing the reversible transfer of CoA groups between acyl substrates. However, how their activities are regulated in response to cellular metabolic states remains unclear. Here we identify a dual regulatory mechanism of a CoA transferase from *Thermus thermophilus*, which is controlled by both interaction with an NAD^+^-sensing accessory protein and protein acetylation. The enzyme associates with an alanine dehydrogenase-like protein that lacks catalytic activity but binds NAD^+^ and NADH. Biochemical analyses revealed that inhibition occurs specifically upon NAD^+^ binding, but not NADH, indicating that the alanine dehydrogenase-like protein senses the intracellular NAD^+^/NADH ratio. Cryo-electron microscopy structures of the complex of CoA transferase and alanine dehydrogenase-like protein reveal the structural basis of this redox-dependent inhibition. Furthermore, acetylation of the CoA transferase alleviates the inhibition. Since the NAD^+^/NADH ratio and acetyl-CoA levels reflect the cellular energy and metabolic states, these findings uncover a previously unrecognized regulatory link through which redox sensing and acetylation cooperate to fine-tune the β-oxidation flux.

## Introduction

Protein lysine acetylation, which was first discovered in histones in the 1960s (Allfrey *et al*, 1964; Phillips, 1963), is now recognized as widespread post-translational modifications in all domains of life (Christensen *et al*, 2019; Narita *et al*, 2019; Soppa, 2010; VanDrisse & Escalante-Semerena, 2019). Since the development of a proteomic analysis combined with the affinity purification of acetylated peptides using immunoprecipitation, called an acetylome analysis, for HeLa cells and mouse liver mitochondria (Kim *et al*, 2006), numerous acetylated proteins have been identified in many living organisms (Kosono *et al*, 2015; Mizuno *et al*, 2016; Yoshida *et al*, 2019; Zhang *et al*, 2013). Protein acetylation has been detected in various metabolic enzymes, suggesting its role in the regulation of metabolism through changes in the activity, stability, and/or interactions with other enzymes/proteins responsible for metabolism. In bacteria, acetylation mostly occurs non-enzymatically using acetyl-phosphate and acetyl-CoA as the donor of an acetyl group (Kuhn *et al*, 2014; Weinert *et al*, 2013). Therefore, although many proteins are acetylated in bacterial cells, limited information is currently available on the effects of non-enzymatic acetylation on the functions of specific proteins. On the other hand, reversible protein acetylation mediated by protein lysine acetyltransferases (KAT) and protein lysine deacetylase (KDAC) is known to play important roles in regulating the activities of several enzymes. AMP-forming acetyl-CoA synthetase (ACS) from *Salmonella enterica* is regulated through reversible protein acetylation by acetyltransferase (Pat/YfiQ) and deacetylase (CobB) in response to acetyl-CoA availability in cells (Starai *et al*, 2002; Starai & Escalante-Semerena, 2004; Starai *et al*, 2004). Several AMP-forming ACS are regulated by protein acetylation in combination with transcriptional regulation or cyclic AMP binding (Burckhardt *et al*, 2020; Gardner *et al*, 2006; Kumari *et al*, 2000; Xu *et al*, 2011; You *et al*, 2017). As other examples, enzymes involved in central metabolism, such as glutamine synthetase (You *et al*, 2016), malate dehydrogenase (Venkat *et al*, 2017), isocitrate dehydrogenase (Venkat *et al*, 2018), and formate dehydrogenase (Zhang *et al*, 2020), were shown to be controlled by reversible lysine acetylation. In protein acetylation, enzymatic acetylation is expected to have direct biological functions.

Proteins identified as KATs in bacteria possess Gcn5-related acetyltransferase (GNAT) motifs (Gardner *et al*., 2006; Kumari *et al*., 2000; Starai & Escalante-Semerena, 2004; Xu *et al*., 2011; Zhang *et al*., 2020). In the present study, we attempted to identify proteins acetylated in a KAT-dependent manner and elucidate the role of acetylation in cellular metabolism. To achieve this, we used *Thermus thermophilus* HB27, an extremely thermophilic bacterium, as the target because it has a small genome and only ten genes encoding proteins with GNAT motifs.

We herein detected a protein that was markedly acetylated by TT_C1711 in *T. thermophilus* and identified it as TT_C1083, annotated as a CoA transferase (CoAT). CoAT is an enzyme that catalyzes the transfer of the CoA moiety from acyl-CoA to short-chain fatty acids (SCFAs) in an ATP-independent manner (Heider, 2001). It is also involved in the utilization of butyrate in *Escherichia coli* (Vanderwinkel *et al*, 1968) and the energy-independent production of SCFAs in intestinal bacteria, such as *Clostridium* species (Scherf & Buckel, 1991; Trachsel *et al*, 2016; Vital *et al*, 2013; Wiesenborn *et al*, 1989). In mammals, succinyl-CoA:3-ketoacid CoA transferase (SCOT) plays an important function in ketone body utilization and its gene expression is regulated in a tissue-dependent manner (Fukao *et al*, 1997). CoAT is therefore both metabolically important and subject to diverse regulatory mechanisms, suggesting that similar regulation may occur in TtCoAT. ACS is regulated by KAT-dependent acetylation in response to acetyl-CoA availability in cells and maintains the homeostasis of acetyl-CoA and acetate. By analogy between CoAT and ACS, both of which are related to acyl-CoA metabolism, we hypothesized that CoAT from *T. thermophilus* (TtCoAT) is also regulated by acetylation; however, direct regulation through acetylation was not observed for TtCoAT. We demonstrated that TtCoAT formed a heterocomplex with an enzymatically inactive alanine dehydrogenase homolog that sensed the NAD^+^/NADH ratio. The CoAT activity of the complex was inhibited by NAD^+^, but not by NADH. The inhibitory effects of NAD^+^ were reduced by the acetylation of TtCoAT. The present results revealed the control of TtCoAT activity through protein-protein interactions and protein acetylation, which may be involved in the flux control of acyl-CoA and energy metabolism coupled with the β-oxidation pathway. Furthermore, the cryo-electron microscopy (cryo-EM) structure of the TtCoAT·ADLP complex provides the structural basis for the inhibitory mechanism of TtCoAT by binding ADLP·NAD^+^. Our study provides a new example of regulatory control in CoAT, suggesting how enzyme activity can be modulated in response to cellular NAD^+^/NADH ratio and acetyl-CoA availability.

## Results

### Identification of CoA transferase as a highly acetylated protein in T. thermophilus HB27

*T. thermophilus* HB27 carries ten genes that encode proteins with GNAT motifs on the genome. To screen for proteins acetylated in a KAT-dependent manner in *T. thermophilus*, we constructed ten knockout mutants (ΔKATs), each with a corresponding gene deletion by replacement with a kanamycin resistance marker, and compared the acetylation profile of the lysate between the wild-type and ΔKAT strains by western blotting using anti-acetyllysine antibodies. The results obtained revealed a highly acetylated protein of approximately 50 kDa (Protein X) in the lysate of the wild type, but not in the Δ*TT_C1711* lysate (Fig. 1A), suggesting that TT_C1711 functioned as a KAT for Protein X. Through chromatography followed by western blotting using anti-acetyllysine antibodies (Supplementary Fig. S1), we partially purified Protein X from wild-type *T. thermophilus* and identified it as TT_C1083, annotated as 4-hydroxybutyrate CoA transferase, by a MALDI-TOF MS analysis with 79% sequence coverage. We also confirmed that the band of the acetylated Protein X disappeared in western blotting for Δ*TT_C1083* (Fig. 1B).

**Figure 1.**
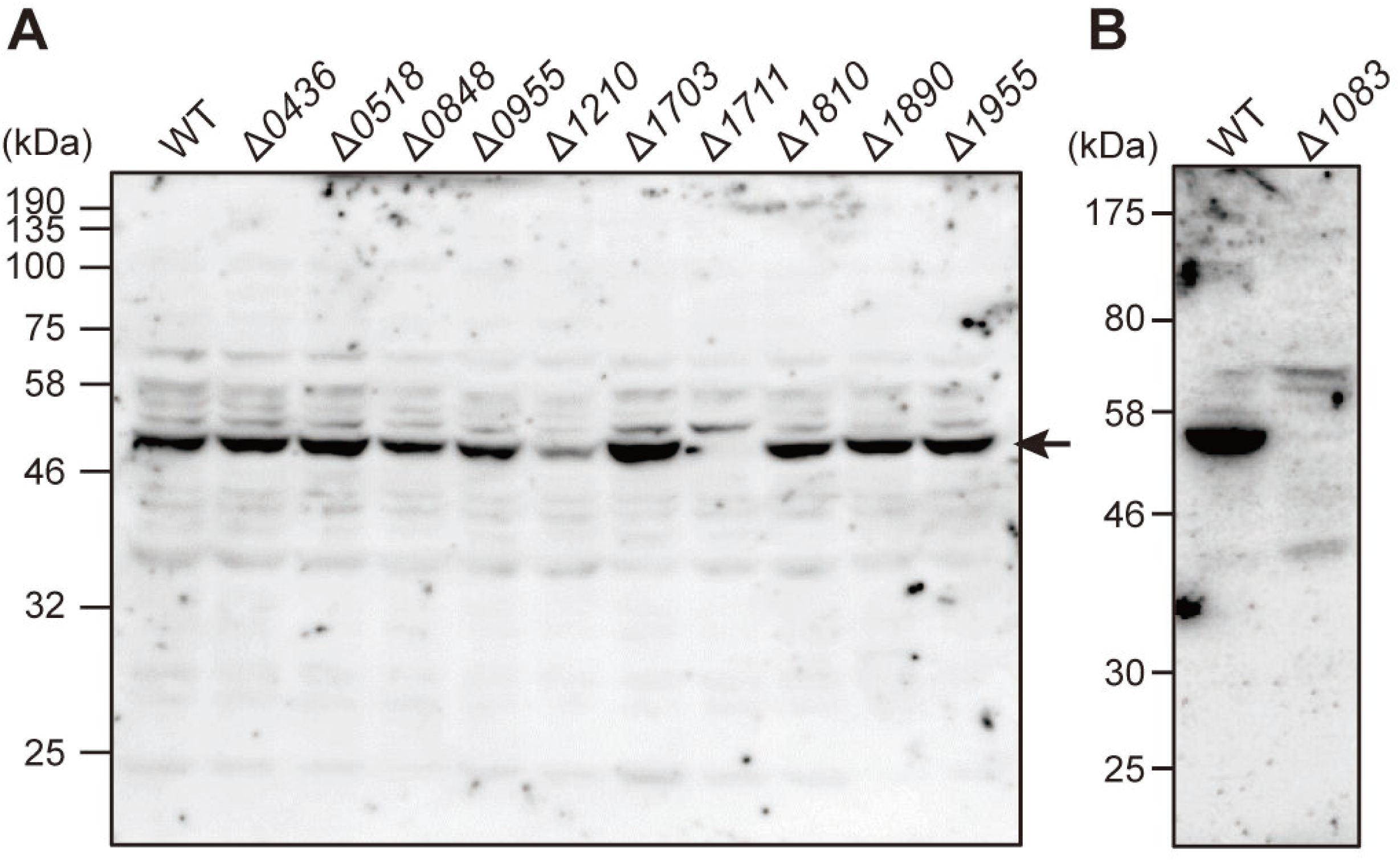
Effects of the knockout of the putative KAT gene on the acetylation profile of *T. thermophilus*. A. Western blotting for the cell extract from the wild-type (lane 1) and disruptants of each KAT candidate gene (lane 2-11) using anti-acetyllysine antibodies. B. Western blotting for the cell extract from the wild type and Δ*TT_C1083* using anti-acetyllysine antibodies. An arrow indicates Protein X (TT_C1083), which was highly acetylated in the wild type.

To confirm that TT_C1711 acetylated TT_C1083 using acetyl-CoA as the donor of an acetyl group, recombinant TT_C1711 and TT_C1083 were prepared and used for the *in vitro* reaction. The results obtained showed that TT_C1711 acetylated TT_C1083 in an acetyl-CoA dependent manner (Fig. 2A). In a subsequent LC-MS/MS analysis, we identified seven lysine residues (Lys5, Lys6, Lys37, Lys357, Lys401, Lys435, and Lys440) of TT_C1083 that were acetylated by TT_C1711. The successive replacement of each lysine residue to alanine using recombinant TT_C1083 confirmed that Lys5 was the main acetylated residue because only the K5A mutant showed a marked decrease in acetylation (Fig. 2B, C). Furthermore, an *E. coli* strain co-producing the TT_C1711 and either TT_C1083 or TT_C1083 K5A proteins was constructed. Western blotting revealed that TT_C1083 co-produced with TT_C1711 was acetylated, whereas TT_C1083 K5A was not even when TT_C1711 was co-produced (Fig. 2D). These results confirmed that Lys5 was the main acetylated residue.

**Figure 2.**
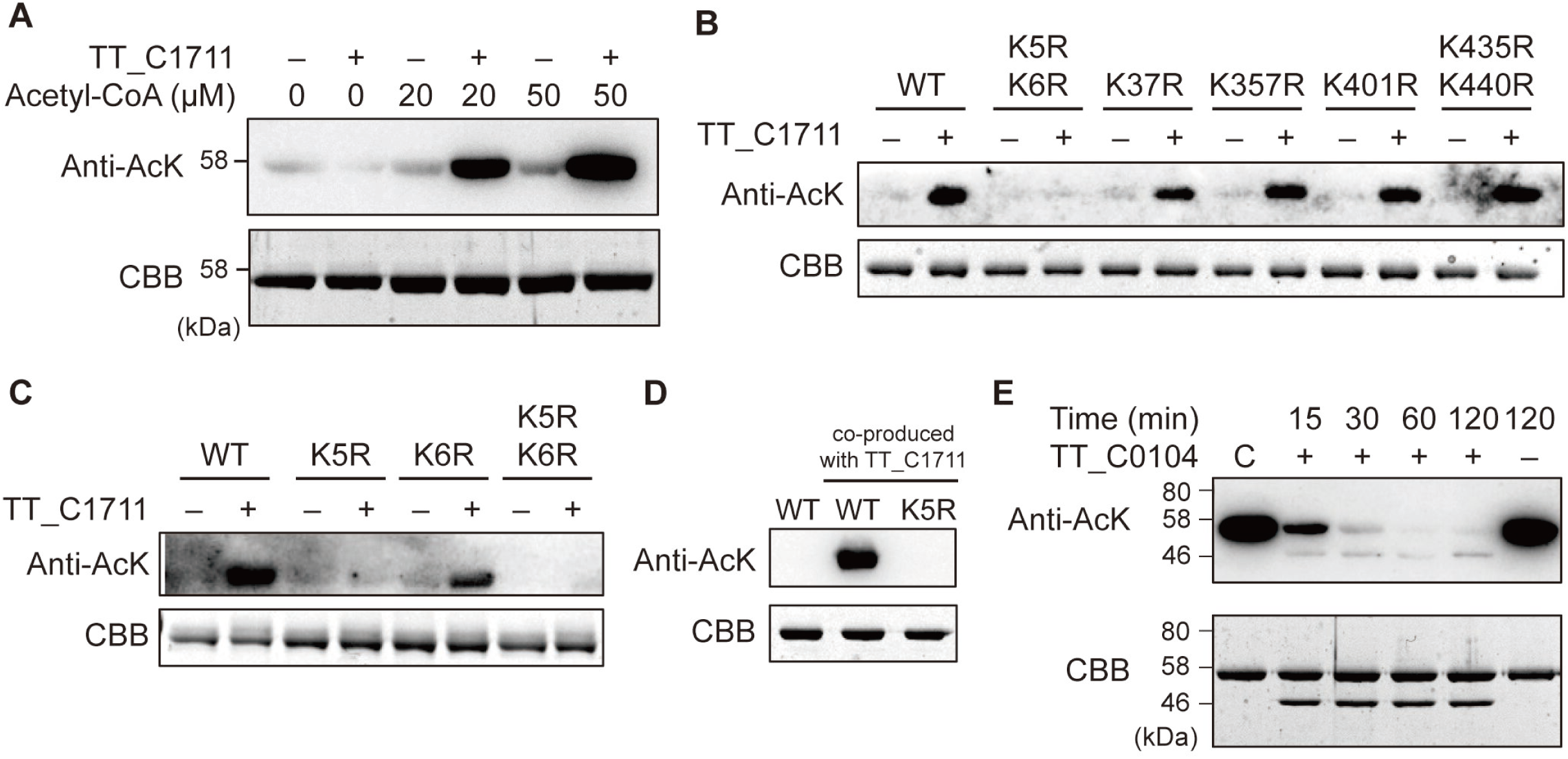
Reversible acetylation of TT_C1083. A. Enzymatic acetylation of TT_C1083 by TT_C1711. B, C. Enzymatic acetylation of TT_C1083 derivatives by TT_C1711. D. *In vitro* acetylation of TT_C1083 by TT_C1711 in *E. coli* cells. The leftmost lane shows TT_C1083 and the remaining lanes indicate TT_C1083 and TT_C1083 K5A co-produced with TT_C1711. E. Deacetylation of acetylated TT_C1083 by TT_C0104. Acetylated TT_C1083 before the deacetylation reaction was applied at the leftmost lane (“C”). The upper and lower panels show western blots using anti-acetyllysine antibodies and Coomassie brilliant blue staining of SDS-PAGE, respectively.

Since the KAT-dependent acetylation of TT_C1083 was expected to affect the function of this protein, a deacetylation system coupled with an acetylation system was anticipated in *T. thermophilus*. In *T. thermophilus*, three genes, *TT_C0104*, *TT_C1026*, and *TT_C1955*, are annotated to encode deacetylases. To identify the KDAC responsible for the deacetylation of TT_C1083, we purified three recombinant deacetylase homologs. An *in vitro* assay revealed that only TT_C0104, a metal-dependent deacetylase, deacetylated TT_C1083 that was already acetylated by TT_C1711 (Fig. 2E and Supplementary Fig. 2A). Although deacetylation was only observed in the absence of ZnSO_4_, the addition of EDTA, which chelates Zn^2+^ ions, to the reaction mixture had a negative effect on the deacetylation of acetylated TT_C1083 (Supplementary Fig. 2B). Moreover, the addition of butyrate, which bound to the Zn^2+^ site of KDAC, also inhibited the deacetylation of acetylated TT_C1083 (Supplementary Fig. 2B). These results suggest that TT_C0104 tightly bound Zn^2+^ ions and an excess concentration of Zn^2+^ ions exerted inhibitory effects on its activity. Collectively, these results indicate the control of the acetylation status of TT_C1083 by the combination of TT_C1711 (KAT) and TT_C0104 (KDAC).

### Characterization of TT_C1083

To elucidate the function of TT_C1083 from *T. thermophilus*, we prepared the recombinant TT_C1083. Since TT_C1083 is annotated as 4-hydroxybutyrate CoA transferase, an enzyme assay was performed using various SCFAs as substrates. The results obtained showed that TT_C1083 exhibited CoA-transferring activity for various SCFAs using acetyl-CoA as the CoA donor (Fig. 3A, B). To confirm the function of TT_C1083 *in vivo*, we constructed a knockout strain of *TT_C1083* (Δ*TT_C1083*).

**Figure 3.**
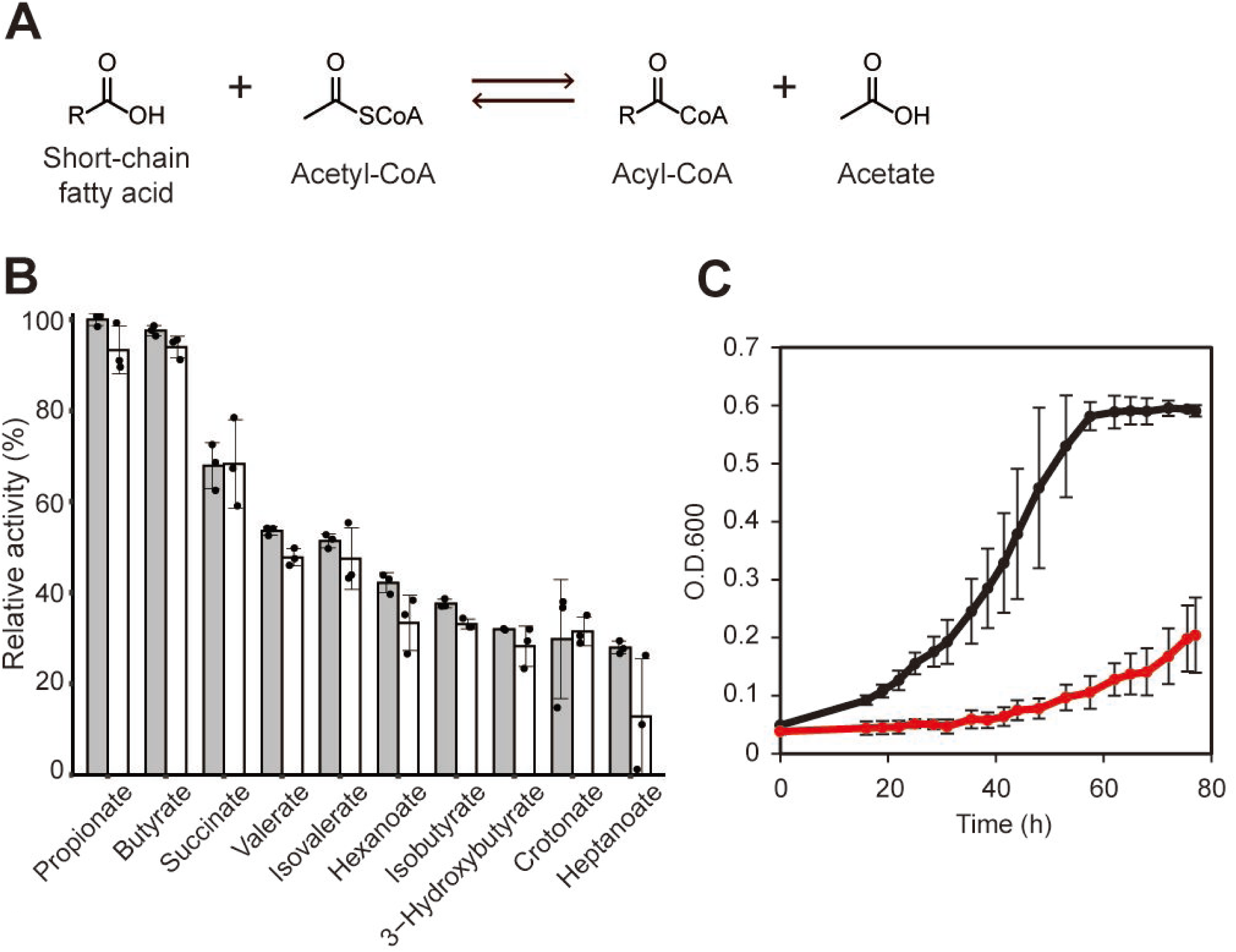
Function of CoAT A. Scheme of the CoAT reaction. B. TtCoAT activities for various substrates. The substrates used are shown under the bar. Gray and white bars indicate the activities of TtCoAT and AcTtCoAT, respectively. Relative activity was shown as the activity of TtCoAT for propionate as 100. C. Growth curves of *T. thermophilus* and the Δ*TtCoAT* strain in minimal medium supplemented with 10 mM butyrate as the sole carbon source. Growth curves shown in black and red lines are of the wild type and Δ*TtCoAT*, respectively. O.D.600, optical density at 600 nm.

Consistent with the *in vitro* activity assay, Δ*TT_C1083* showed a clear growth defect in the minimal medium containing butyrate as the sole carbon source (Fig. 3C). These results indicate that TT_C1083 functioned as an acyl-CoA transferase and was involved in the utilization of SCFAs, such as butyrate; therefore, it was hereafter referred to as TT_C1083 TtCoAT.

TtCoAT was identified as a major acetylated protein in *T. thermophilus*. TtCoAT converts SCFAs to acyl-CoAs, which are metabolized to acetyl-CoA, and, thus, it plays a role in acetyl-CoA recycling/synthesis, similar to ACS. ACS is known to be regulated by protein acetylation in response to acetyl-CoA concentrations in cells. Since ACS and CoAT both use acetyl-CoA as a substrate and are acetylated in cells, we hypothesized that CoAT may be regulated by protein acetylation. TtCoAT was produced in a highly acetylated form (AcTtCoAT) when it was co-produced with TT_C1711 in *E. coli* cells. However, when we prepared TtAcCoAT and then compared its CoAT activity with that of non-acetylated TtCoAT, no significant differences were observed in CoAT activity or its substrate specificity by protein acetylation (Fig. 3B).

### CoAT interacts with an enzymatically inactive alanine dehydrogenase homolog

We postulated that acetylation effectively regulates TtCoAT indirectly through interactions with other (regulatory) proteins. To confirm this hypothesis, we screened a partner protein of TtCoAT that interacted with TtCoAT using *T. thermophilus* producing TtCoAT with an affinity tag. In sodium dodecyl sulfate-polyacrylamide gel electrophoresis (SDS-PAGE) of the concentrated eluate, we observed a protein of 40 kDa was co-purified with TtCoAT (Fig. 4A). This protein was identified by in-gel digestion followed by a MALDI-TOF-MS analysis as TT_C1082, annotated as an alanine dehydrogenase. A pull-down assay using recombinant TtCoAT with a (His)_6_-tag and TT_C1082 with a Strep-tag confirmed that both proteins were co-purified using Ni^2+^-NTA or Strep-tactin resins (Fig. 4B, C). When the co-purified proteins were applied to size exclusion column chromatography, both proteins were eluted as a single peak corresponding to 401 kDa, indicating that they form a stable complex. (Fig. 4D). Since TtCoAT or ADLP alone forms a tetramer (159 kDa) and hexamer (186 kDa), respectively, the protein complex detected was estimated to be a hetero-decamer composed of the TtCoAT tetramer and ADLP hexamer.

**Figure 4.**
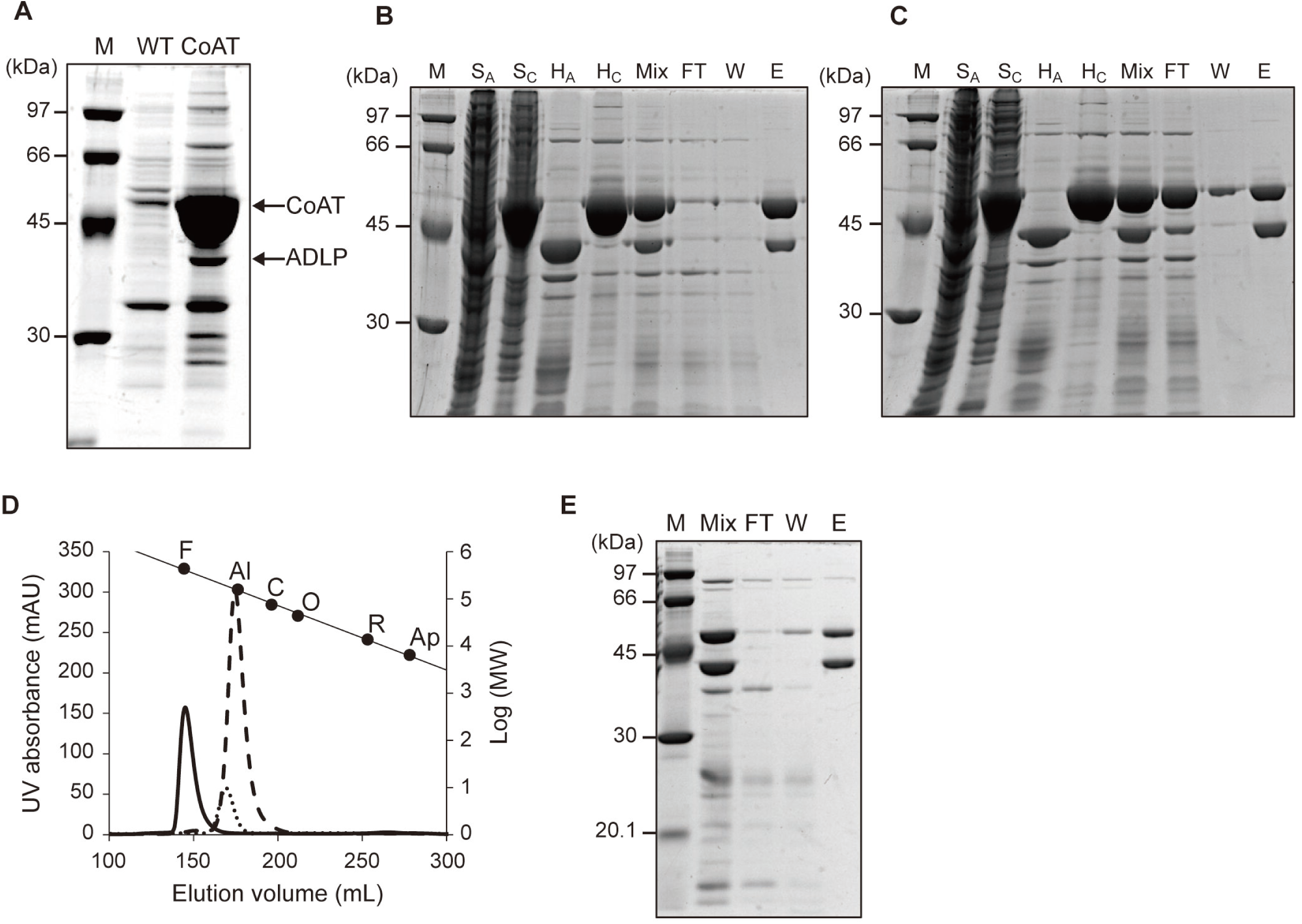
TtCoAT interacts with ADLP. A. Pull-down assay for Strep-tagged TtCoAT using a Strep-tactin column. Lane M, molecular markers; lane WT, elution from the wild-type strain; lane CoAT, elution from TtNStrepHisCoAT. B, C. Co-purification of TtCoAT with the His-tag at the C terminus and ADLP with Strep-tag at the N terminus. SDS-PAGE of fractions from the purification steps is shown. Ni^2+^-NTA resin and Strep-tactin resin are used as the affinity purification steps for (B) and (C), respectively. Lane M, molecular markers; lane S_A_ and S_C_, supernatants from *E. coli* producing ADLPNstrep and TtCoATCHis, respectively; lane H_A_ and H_C_, samples after the heat treatment of ADLPNstrep and TtCoATCHis, respectively; lane Mix, Mix of H_A_ and H_C_ samples; lane FT, flow-through fraction; lane W, washing fraction; lane E, eluate fraction. D. Elution profiles of TtCoAT, ADLP, and the TtCoAT·ADLP complex in gel-filtration chromatography. UV absorbance at 280 nm for TtCoAT, ADLP, and the TtCoAT·ADLP complex is shown in dashed, dotted, and solid lines, respectively. Molecular size markers are plotted for their elution volumes. F, Ferritin (440 kDa); Al, Aldolase (158 kDa); C, Conalbumin (75 kDa); O, Ovalbumin (44 kDa); R, Ribonuclease A (13.7 kDa); Ap, Aprotinin (6.5 kDa). E. SDS-PAGE of AcTtCoAT (with His-tag) and ADLP (with Strep-tag), which were separately purified and mixed *in vitro*. Strep-tactin resin was used for the purification step. Lane M, molecular markers; lane Mix, Mix of AcTtCoAT and ADLP partially purified by the heat treatment; lane FT, flow-through fraction; lane W, washing fraction; lane E, eluate fraction.

*T. thermophilus* HB27 possesses another alanine dehydrogenase gene, *TT_C1770*, on its genome. We also prepared recombinant TT_C1770 from *E. coli* and analyzed the activities of both alanine dehydrogenase homologs. TT_C1770 exhibited alanine dehydrogenase activity in both reactions, oxidative deamination and reductive amination for alanine and pyruvate, respectively. On the other hand, TT_C1082 showed no activity for alanine or pyruvate (Fig. 5A). We also investigated the activity of TT_C1082 for the other 19 proteinogenic amino acids; however, TT_C1082 did not exhibit any activities for the amino acids tested. Therefore, we named it an <u>a</u>lanine <u>d</u>ehydrogenase-like <u>p</u>rotein, ADLP. Amino acid sequence alignment between ADLP and a typical alanine dehydrogenase revealed that the catalytic lysine residue was replaced by Ala (Ala77) in ADLP, which accounted for the loss of the enzyme activity of ADLP (Supplementary Fig. 3).

**Figure 5.**
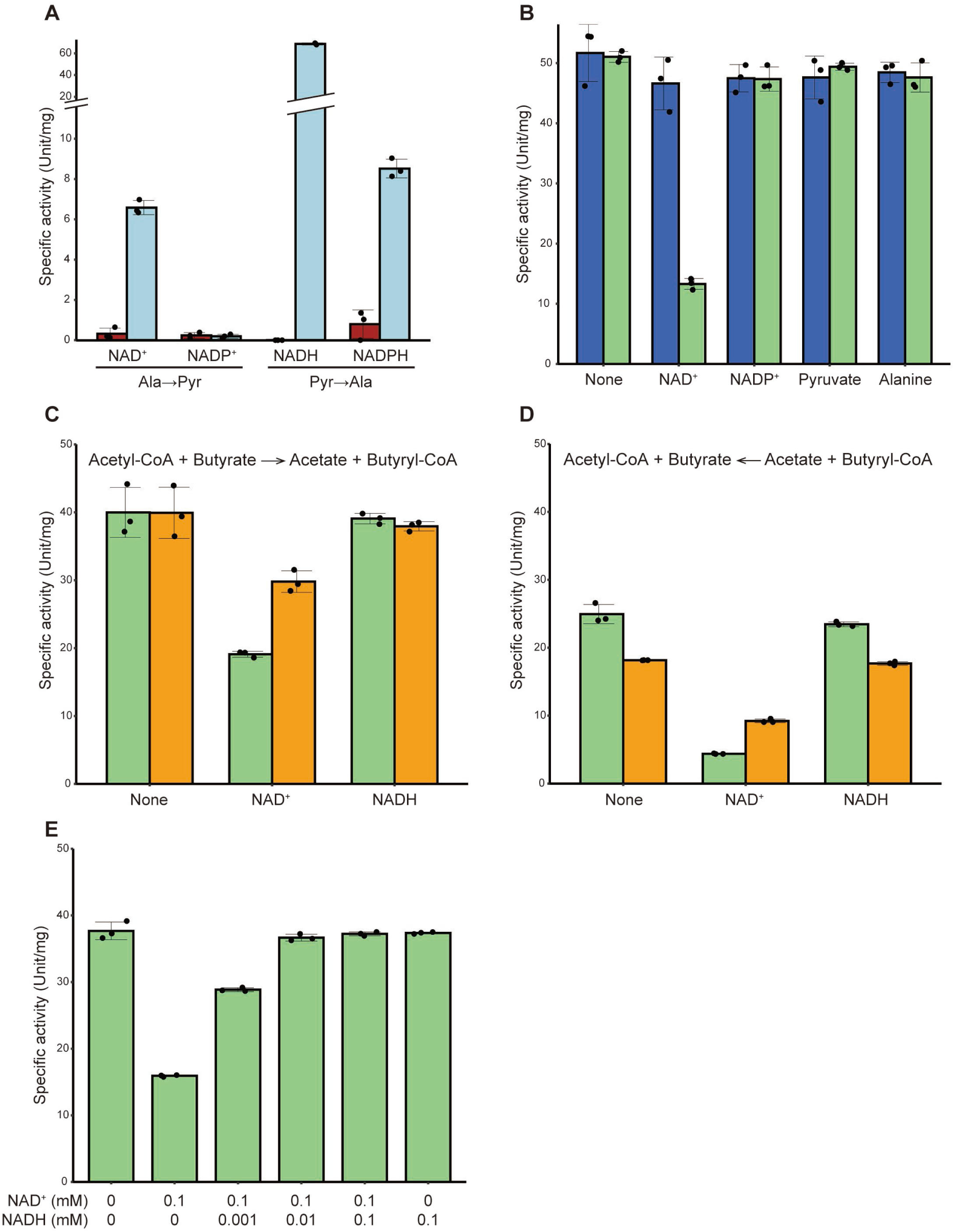
ADLP is an enzymatically inactive regulatory protein for TtCoAT. A. Alanine dehydrogenase activities of AlaDH and ADLP from *T. thermophilus*. The specific activities of ADLP and AlaDH are shown in red and light blue bars, respectively. B. CoAT activity in the presence of ADLP and/or NAD(P)^+^, alanine, or pyruvate. The specific activities of TtCoAT in the presence or absence of ADLP are shown in blue and light green bars, respectively. C, D. CoAT activity in the presence of ADLP and NAD(H). The specific activities of TtCoAT·ADLP and AcTtCoAT·ADLP are shown in light green and orange bars, respectively. C shows the butyryl-CoA synthesis reaction, and D shows the acetyl-CoA synthesis reaction. E. CoAT activity of the TtCoAT·ADLP complex in the presence of various concentrations of NAD^+^ and NADH.

### ADLP binds NAD^+^ to control CoAT activity

ADLP lacked the catalytic lysine residue of alanine dehydrogenase, as described above, but retained most of the residues responsible for substrate and NAD(H) binding (Supplementary Fig. 3). Therefore, we hypothesized that ADLP serves as a regulatory protein of TtCoAT to bind alanine, pyruvate, and/or NAD(P)(H). To confirm this hypothesis, the effects of ADLP and possible effector compounds on CoAT activity were examined using butyryl-CoA and acetate as the substrates in a coupling reaction with a citrate synthase (CS). The results obtained showed that CoAT activity decreased when ADLP and NAD^+^ were both present, whereas NAD^+^ alone did not affect CoAT activity without ADLP (Fig. 5B). We also directly detected the reaction product, acyl-CoAs, in the presence or absence of ADLP and NAD(H) and examined the effects of NADH, which interfered with the above coupling system. The assay confirmed the inhibition of TtCoAT by NAD^+^ in the presence of ADLP; 48% inhibition in the conversion of acetyl-CoA and butyrate to acetate (Fig. 5C) and butyryl-CoA and 18% inhibition in the reverse reaction (Fig. 5D). On the other hand, NADH did not affect CoAT activity in the presence or absence of ADLP. These results suggest that NAD^+^-bound ADLP interacted with TtCoAT to negatively control CoAT activity.

To investigate whether the TtCoAT·ADLP complex binds NADH as well as NAD^+^, the binding of these coenzymes to the complex was investigated by isothermal titration calorimetry (ITC). The results obtained indicated that the TtCoAT·ADLP complex bound NAD^+^ and NADH. The binding affinity (*K*_d_) of the TtCoAT·ADLP complex for NADH was markedly higher than that for NAD^+^ (*K*_d_^NAD+^ = 3.1 μM, *K*_d_^NADH^ = 0.34 μM) (Table 1). We also confirmed that NAD^+^ and NADH both bound to ADLP with similar affinities (*K*_d_^NAD+^ = 9.1 μM, *K*_d_^NADH^ = 0.22 μM) in the absence of CoAT (Supplementary Table S1). Consistent with the difference in the affinity of the TtCoAT·ADLP complex to NAD^+^ and NADH, the inhibitory effects of NAD^+^ on CoAT activity were canceled by the addition of NADH at a concentration of 1/10 of NAD^+^ (Fig. 5E). Moreover, additional ITC measurements indicated that ADLP did not bind acetate, butyrate, acetyl-CoA, or butyryl-CoA, which were used in the activity assay of TtCoAT (Supplementary Fig. S4). These results imply that ADLP was involved in controlling CoAT activity by forming the TtCoAT·ADLP complex in response to the NAD^+^/NADH ratio in cells.

**Table 1.** Thermodynamic parameters of NAD(H) binding to the TtCoAT·ADLP complex

|  | NAD<sup>+</sup> | NADH |
| --- | --- | --- |
| N (sites) | 3.0 ± 0.05 | 3.2 ± 0.03 |
| *K<sub>d</sub>* (μM) | 3.1 ± 0.33 | 0.34 ± 0.051 |
| Δ*H* (kcal/mol) | -18.7 ± 0.5 | -15.5 ± 0.2 |
| Δ*S* (cal/mol/deg) | -36.3 | -21.5 |

### The TtCoAT and ADLP interaction may change in the presence of NAD^+^

The mechanisms by which the binding of NAD^+^, but not NADH, to ADLP affects TtCoAT activity need to be elucidated in more detail. Therefore, we investigated the affinity between TtCoAT and ADLP in the presence or absence of NAD^+^ and NADH using ITC. In the absence of the effectors, the *K*_d_ of the ADLP hexamer to the TtCoAT tetramer was calculated as 0.81 μM (Table 2). This thermodynamic parameter showed that the interaction between TtCoAT and ADLP was driven by enthalpy. The *K*_d_ value showed an approximately 20-fold reduction (34 nM) in the presence of 1 mM NAD^+^. Conversely, 1 mM NADH did not affect *K*_d_ (*K*_d_ = 0.53 μM). These results indicate that the binding of NAD^+^ to ADLP stabilized the interaction between TtCoAT and ADLP, resulting in the inhibition of CoAT.

**Table 2.**
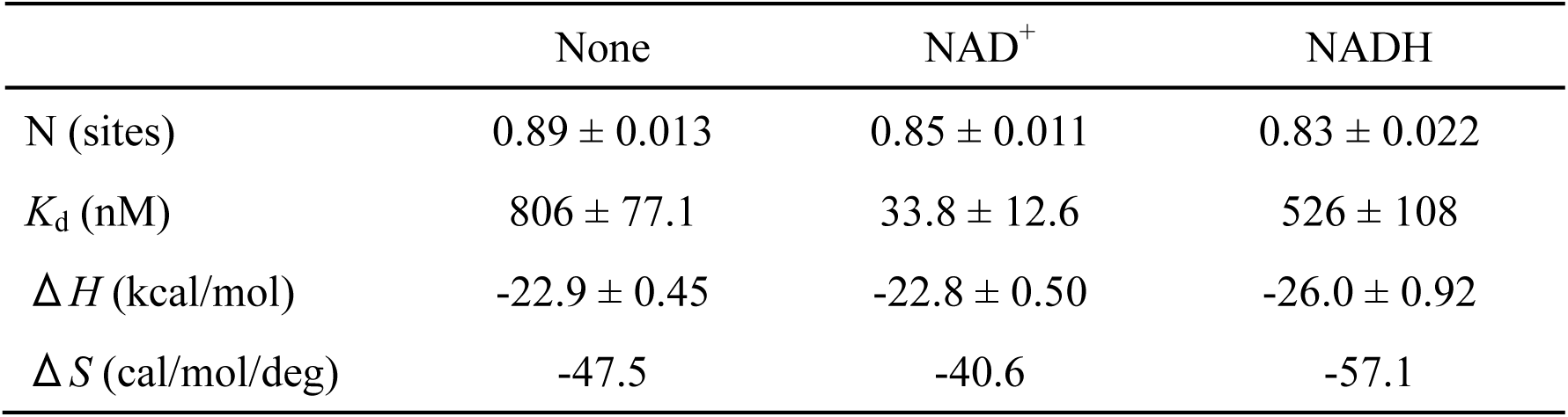
Thermodynamic parameters of the ADLP hexamer binding to the TtCoAT tetramer

### TtCoAT acetylation partially alleviates the inhibition by ADLP·NAD^+^

As described above, the KAT-dependent acetylation of TtCoAT did not affect the activity of CoAT in the absence of ADLP. To elucidate the roles of acetylation on CoAT activity, we examined the effects of the protein acetylation of CoAT on the interaction between CoAT and ADLP and/or the inhibition of CoAT activity by ADLP·NAD^+^. A pull-down assay showed that ADLP was co-purified with AcTtCoAT (Fig. 4E), suggesting that acetylation did not affect the protein-protein interaction. On the other hand, as described above, 1 mM NAD^+^ decreased CoAT activity in the acetyl-CoA-producing reaction to 18% of that for TtCoAT·ADLP. The same concentration of NAD^+^ reduced the inhibition (50%) of AcTtCoAT·ADLP (Fig. 5D). A similar acetylation-mediated reduction in inhibition was observed in the reverse reaction to produce butyryl-CoA (48 to 78 %) (Fig. 5C). These results imply that the protein acetylation of TtCoAT partially canceled the inhibition by ADLP·NAD^+^. To elucidate the effects of acetylation and NAD^+^ on the catalytic properties of TtCoAT, we performed a kinetic analysis of TtCoAT·ADLP and AcTtCoAT·ADLP using butyryl-CoA and acetate as the substrates. As expected, the apparent kinetic parameters were similar between TtCoAT·ADLP and AcTtCoAT·ADLP, giving the same catalytic efficiency, *V*_max_/*K*_m_ ^butyryl-CoA^, of 1.85 units·mg^-1^·μM^-1^ in the absence of NAD^+^. In the presence of NAD^+^, the *V*_max_ values of TtCoAT·ADLP and AcTtCoAT·ADLP both showed approximately 3-fold decreases. On the other hand, a marked difference was observed in the apparent *K*_m_ value for butyryl-CoA. NAD^+^ did not affect the apparent *K*_m_ value for the butyryl-CoA of AcTtCoAT·ADLP, but improved the *K*_m_ value of TtCoAT·ADLP by approximately 1.6-fold (Table 3). These results showed the regulation of CoAT activity in *T. thermophilus* by both an interaction with the regulatory protein (ADLP) that senses the NAD^+^/NADH ratio and protein acetylation.

**Table 3.** Kinetics parameters of TtCoAT·ADLP in the presence and absence of NAD^+^.

| Enzyme | NAD<sup>+</sup> (mM) | *V<sub>max</sub>* (unit/mg) | *K<sub>m</sub>*<sup>*</sup> (μM) | *V<sub>max</sub>*/*K<sub>m</sub>*<sup>*</sup> (unit/mg/μM) |
| --- | --- | --- | --- | --- |
| TtCoAT·ADLP | 0 | 61.4 ± 1.9 | 33.1 ± 2.8 | 1.85 ± 0.17 |
|  | 0.1 | 17.0 ± 1.2 | 31.0 ± 6.3 | 0.55 ± 0.12 |
| AcTtCoAT·ADLP | 0 | 56.9 ± 2.0 | 30.7 ± 3.0 | 1.85 ± 0.19 |
|  | 0.1 | 20.7 ± 0.9 | 19.6 ± 3.0 | 1.06 ± 0.17 |

\* Apparent *K<sub>m</sub>* value for butyryl-CoA

### Cryo-EM structures of the TtCoAT·ADLP complex

To clarify the molecular mechanisms underlying the regulation of CoAT, we successfully determined the structure of the TtCoAT·ADLP complex in the NAD^+^-bound and NAD^+^-free forms at 2.3 and 2.7 Å, respectively, using cryo-EM (Fig. 6A, B and Supplementary Fig. S5). The NAD^+^-bound form of the TtCoAT·ADLP complex consisted of six chains (A-F chains) of ADLP and six chains of TtCoAT (G-L chains) (Fig. 6A). Most of the ADLP residues were modeled; however, the residues in the I and J chains of TtCoAT were only partially modeled. The NAD^+^-free form of the TtCoAT·ADLP complex was modeled to contain six ADLP subunits (A-F chains) and two or a few TtCoAT subunits (G-J chains) (Fig. 6B), in which most of the residues in the G and H chains were modeled, but only some were observed for the I and J chains. Several regions in each ADLP subunit in the NAD^+^-free complex were not modeled, particularly loops 240-247 and 270-280 in the nucleotide-binding domain responsible for NAD^+^ binding, which may have been due to the flexibility of these loops that did not bind NAD^+^. The structure of ADLP was similar to that of typical AlaDH, such as AlaDH from *Mycobacterium tuberculosis* (MtAlaDH), forming a hexamer composed of a trimer of dimers (Agren *et al*, 2008; Tripathi & Ramachandran, 2008) (Supplementary Fig. S6A, B). TtCoAT also formed dimers as its structural unit, adopting a similar structure to that of known class I CoATs, such as 4-hydroxybutyrate CoA transferase from *Clostridium aminobutyricum* (Macieira *et al*, 2012; Macieira *et al*, 2009) (Supplementary Fig. S7A, B). Both structures of the TtCoAT·ADLP complex had an ADLP hexamer in the center and one to three dimers of TtCoAT dimers bound around it. Based on the results of gel filtration, the complex was initially predicted to consist of a CoAT tetramer and ADLP hexamer; however, the eluted proteins were likely in a state of equilibrium, in which one to three CoAT dimers bound to an ADLP hexamer. The most prominent difference between the NAD^+^-bound and NAD^+^-free forms of the TtCoAT·ADLP structures was the number of TtCoAT dimers interacting with the ADLP hexamer (Fig. 6A, B). In addition to the clear electron densities for the two TtCoAT dimers (GH and KL chains) in the NAD^+^-bound form, weak electron densities for the third TtCoAT dimer (IJ chains) were observed. On the other hand, in the NAD^+^-free form, only one molecule of the TtCoAT dimer with clear electron densities interacted with the ADLP hexamer. This is consistent with the difference in affinity between CoAT and ADLP in the presence and absence of NAD^+^, as revealed by the ITC analysis described above.

**Figure 6.**
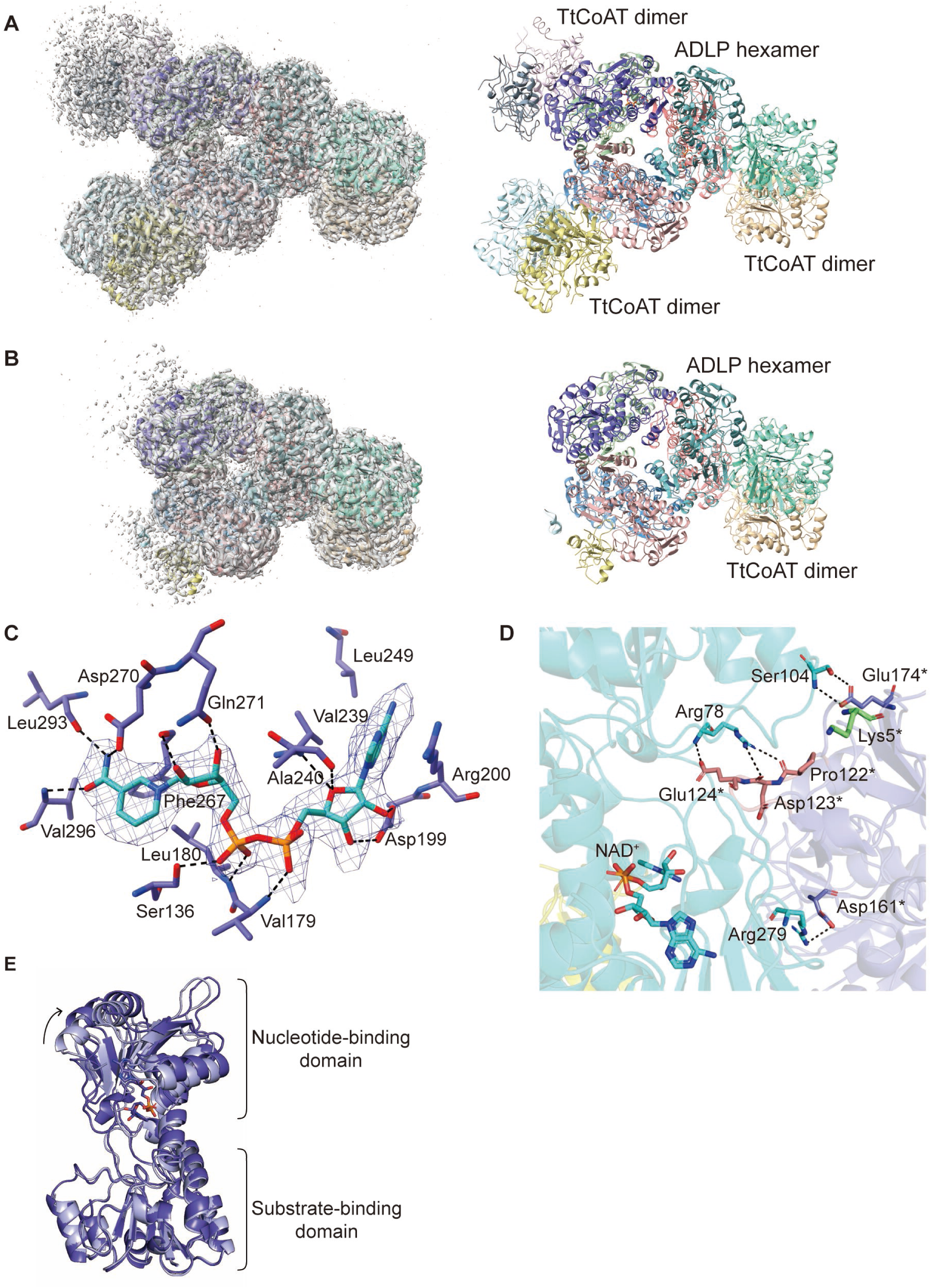
Cryo-EM structures of the TtCoAT·ADLP complex. A, B. Cryo-EM density maps (left) and corresponding models (right) of the TtCoAT·ADLP complex in the NAD^+^-bound form (A) and NAD^+^-free form (B). C. Close-up view of the NAD^+^-binding site in ADLP with a density map of NAD^+^ and the surrounding residues. NAD^+^ is shown as a stick model in cyan, and interacting residues are depicted as stick models in blue. D. Close-up view of the interaction interface between chain B and chain G. Loop 120-126 of chain G in TtCoAT, inserted into the cleft of ADLP, is highlighted in pink. Lys5, which is an acetylated site of TtCoAT, is shown in pale green. Residues involved in the interaction between TtCoAT and ADLP are shown in stick models. E. Conformational change in ADLP upon NAD^+^ binding. The structures of ADLP in the NAD^+^-free form (chain A, pale cyan) and ADLP in NAD^+^-bound form (chain A, blue) are superimposed at the substrate-binding domain (residues 1-129 and 300-340).

In the NAD^+^-bound form, the electron densities of NAD^+^ were observed in each subunit of the ADLP hexamer. NAD^+^ was recognized by several residues in a similar manner to MtAlaDH (Fig. 6C, Supplementary Fig. S6C, D). The adenine ring of NAD^+^ is recognized by hydrophobic interactions with Arg200, Val239, and Leu249. The 2′- and 3′-OH groups of the adenine ribose form hydrogen bonds with the side chain of Asp199 and the oxygen atom of the adenine ribose is stabilized by hydrogen bonds that form with the α-amino group and the main chain oxygen atom of Ala240. The diphosphate group of NAD^+^ interacts via hydrogen bonds with the main-chain nitrogen atoms from Val179-Leu180 and the side chain of Ser136. The side chains of Asp270 and Gln271 form hydrogen bonds with the 2′- and 3′-OH groups of the nicotinamide ribose, and the nicotinamide ring is recognized through the main chain atoms of Phe267, Leu293, and Val296 and the side chain atom of Asp270 by hydrogen bonds.

In the interaction between ADLP and TtCoAT dimers, each subunit of the TtCoAT dimer interacted with each subunit of the ADLP dimer in a different manner (Supplementary Fig. S8A). For example, loop 120-126 of TtCoAT chain G was inserted into the cleft between the substrate-binding domain and the nucleotide-binding domain of ADLP chain B (Fig. 6D). Asp123-His125 of TtCoAT chain H interacted with the substrate-binding domain of ADLP chain D, specifically with Glu2, Arg69, and Arg88 (Supplementary Fig. S8B). We then compared the structures of ADLP between the NAD^+^-bound and NAD^+^-free forms. When each monomer from the ADLP hexamer was superimposed at the substrate-binding domain of ADLP, the relative position of the nucleotide-binding domain was found to be more open in the NAD^+^-bound form (Fig. 6E). These results indicate that the binding of NAD^+^ stabilized the open conformation of ADLP, which facilitated the insertion of loop 120-126 of TtCoAT into the cleft of ADLP, thereby promoting the formation of a stable TtCoAT·ADLP complex.

The active site of TtCoAT was not involved in the interaction with ADLP, but was facing outward from the complex, allowing potential access by substrates (Supplementary Fig. S5). We then compared the structures of TtCoAT between the NAD^+^-bound and NAD^+^-free complexes, and found no significant structural differences (the root mean square deviation between the GH chain dimers of TtCoAT in the NAD^+^-bound and NAD^+^-free complexes was 0.50 Å). This suggests that the conformation of TtCoAT was fixed when it is bound to ADLP. In the reaction cycle of CoAT, conformational changes in the loops around the active site, along with domain motions between the N- and C-terminal domains, are required for substrate binding and product release (Fraser *et al*, 2010; Macieira *et al*., 2012; Mullins & Kappock, 2012; Torres *et al*, 2011). In the TtCoAT·ADLP complex, the flexible G(V/I)G loop (Gly211-Ile212-Gly213 in TtCoAT), which is known to undergo conformational changes during the catalytic cycle, adopted an open conformation. This result implies that TtCoAT had difficulty adopting the closed conformation required to stabilize the enzyme-acyl intermediate (Supplementary Fig. S7C). A previous study reported that the interaction between subunits within the CoAT dimer changed upon substrate binding (Murphy *et al*, 2016). Therefore, the interaction between the ADLP dimer and TtCoAT dimer may restrict the structural flexibility of TtCoAT, resulting in the inhibition of TtCoAT. Although the acetylation site of TtCoAT, Lys5, was not directly involved in the interaction with ADLP, the N-terminal tail including Lys5 was closed to the substrate-binding domain of the ADLP subunit interacting with loop 120-126 from TtCoAT (Fig. 6D). The acetylation of Lys5 was expected to induce subtle conformational changes in the N-terminal tail and decrease the interaction between TtCoAT and ADLP, thereby restoring the structural flexibility of TtCoAT and alleviating its inhibition.

## Discussion

### Catalytically inactive pseudoenzymes involved in the regulation of metabolic enzymes in T. thermophilus

We revealed that TtCoAT interacted with ADLP, which shows amino acid sequence similarities to alanine dehydrogenase, but is enzymatically inactive, possibly due to the replacement of the catalytic lysine residue at the active site. ADLP retained the ability to bind NAD(H) in order to function as a regulatory protein for TtCoAT in response to the change in the NAD^+^/NADH ratio in cells. In *T. thermophilus*, the genes encoding TtCoAT and ADLP are located next to each other (*TT_C1083* and *TT_C1082*, respectively) and are presumably transcribed as a single transcription unit. This gene cluster for CoAT and ADLP is found in other *Thermus* species and several bacteria, such as *Ignavibacterium album*, *Melioribacter roseus*, and *Roseiflexus* species, suggesting that the ADLP/NAD^+^-mediated regulatory system for CoAT is conserved in *Thermus* species and in some bacteria.

The phylogenetic analysis suggested the clade containing ADLP was branched and clustered separately from that of the canonical alanine dehydrogenases including TT_C1770 (Fig. 5F). The catalytic lysine residue (Lys73 in TT_C1770) was replaced with other amino acids, such as alanine, serine, glycine, aspartate, and threonine, in all the proteins in the ADLP clade, implying that these ADLP homologs do not function as alanine dehydrogenase, but as a regulatory protein of CoAT. The distinct similarity (45.7%) in the amino acid sequence of ADLP to that of alanine dehydrogenase (TT_C1770) suggests that ADLP evolved from the ancestral alanine dehydrogenase through gene duplication and lost catalytic activity while maintaining the ability to bind NAD(H). We previously demonstrated that glutamate dehydrogenase (GDH) was regulated by the catalytically inactive GDH homolog for activation by leucine in *T. thermophilus* (Tomita *et al*, 2011; Tomita *et al*, 2010). In contrast to typical GDH, the GDH of *T. thermophilus* is a hetero-oligomer composed of two similar subunits, GdhA and GdhB, which serve as the catalytically inactive regulatory subunit and catalytic subunit, respectively. We also recently showed that an adenine phosphoribosyltransferase homolog (APRTh) without enzymatic activity formed a complex with the GdhAB hetero-oligomer to enhance GDH activity by binding AMP (Tomita *et al*, 2019). The complex regulatory system of GDH using two enzyme homologs lacking catalytic activities (pseudoenzymes) suggests the importance of regulating GDH to control the balance of carbon-nitrogen metabolism and energy levels in cells. The present study has provided a third example using pseudoenzymes for enzyme regulation in *T. thermophilus*. We assumed that the use of enzyme homologs lacking their original function, but retaining their ability to form an oligomeric complex or to bind their substrate/product, may have been advantageous for the development of a complex regulatory mechanism for metabolic enzymes, particularly in *T. thermophilus* living under thermal conditions.

The acetylation of TtCoAT alleviates the inhibition by ADLP·NAD^+^. Mass spectrometry and mutational analyses revealed that Lys5 at the N terminus was the main acetylated lysine in TtCoAT. Several crystal structures of CoAT homologs, such as 4-hydroxybutyrate CoA transferases from *C. aminobutyricum* and *Yersinia pestis*, are available as the complex with CoA (Macieira *et al*., 2009; Torres *et al*., 2011). Although Lys5 of TtCoAT is not conserved in those CoAT homologs, the N-terminal residues are located far from their catalytic centers in their structures, accounting for no change in its activity by Lys5 acetylation itself in the absence of ADLP. We speculate that Lys5 may be located near the site for interacting with ADLP and slightly change the interaction between TtCoAT and NAD^+^-bound ADLP upon acetylation. However, Lys5 is only conserved in CoAT from *Thermus* species among organisms encoding CoAT and ADLP adjacent to each other on the genomes, suggesting that the modulation of CoAT inhibition by acetylation is a characteristic feature limited to *Thermus* species.

### Physiological impact of the unique regulation system of CoAT

The growth defect of Δ*TtCoAT* in minimal media with butyrate as the sole carbon source suggested that TtCoAT functioned in butyrate utilization *in vivo*. However, the necessity of such a complex regulatory mechanism of TtCoAT only for butyrate utilization by *T. thermophilus* remains unclear. We currently assume that this regulation was developed for homeostasis in cells by controlling acetyl-CoA and energy levels in β-oxidation. In the β-oxidation of longer fatty acids, the butyryl-CoA formed is further metabolized into two molecules of acetyl-CoA. TtCoAT also catalyzes the reverse reaction, the conversion of butyryl-CoA and acetate to equimolar amounts of butyrate and acetyl-CoA (Fig. 7). Therefore, the CoAT reaction may compete with the β-oxidation pathway by branching to yield the corresponding SCFAs. The β-oxidation pathway may produce two acetyl-CoA from one butyryl-CoA, which is more efficient than the CoAT reaction producing one acetyl-CoA. Moreover, the β-oxidation of butyryl-CoA may produce NADH, which is used for ATP synthesis through oxidative phosphorylation. Based on these backgrounds, we hypothesize the physiological role of the regulation of TtCoAT by the interaction with the NAD(H)-binding regulatory protein and protein acetylation as follows. When the NAD^+^/NADH ratio in cells is low, the conversion of acyl-CoA to acetyl-CoA mediated by the CoAT reaction without the excessive production of NADH is acceptable. On the other hand, under conditions where cells require the efficient synthesis of NADH by increasing the flux of acyl-CoA towards β-oxidation, namely, under conditions where the NAD^+^/NADH ratio is sufficiently high, ADLP binds NAD^+^ to inhibit TtCoAT. Since this inhibition is reversed by NADH binding to ADLP, the bifurcation of acyl-CoA flux to the CoAT reaction or β-oxidation may be controlled in response to the NAD^+^/NADH ratio in cells. Meanwhile, protein acetylation occurs when acetyl-CoA is abundant in cells. In this environment, the acetylation of TtCoAT induces the partial escape of TtCoAT from the inhibition by ADLP·NAD^+^ because it is not necessary to produce acetyl-CoA efficiently by β-oxidation. Therefore, we hypothesize that TtCoAT is regulated in a complex manner to maintain cell homeostasis by controlling the acetyl-CoA pool and energy levels in response to the availability of acetyl-CoA and the redox status in cells. A previous study reported that the intracellular concentration of NAD^+^ (2.6 mM) was markedly higher than that of NADH (83 μM) in glucose-fed, exponentially growing *E. coli* (Bennett *et al*, 2009). If this is the case for *T. thermophilus* cells, CoAT activity may be inhibited in actively growing *T. thermophilus* cells, in which the NAD^+^ level is markedly higher than that of NADH, resulting in effective energy production. Further studies are needed to clarify the relationship between the function of TtCoAT, the redox status, acyl-CoA and SCFA pools, and protein acetylation.

**Figure 7.**
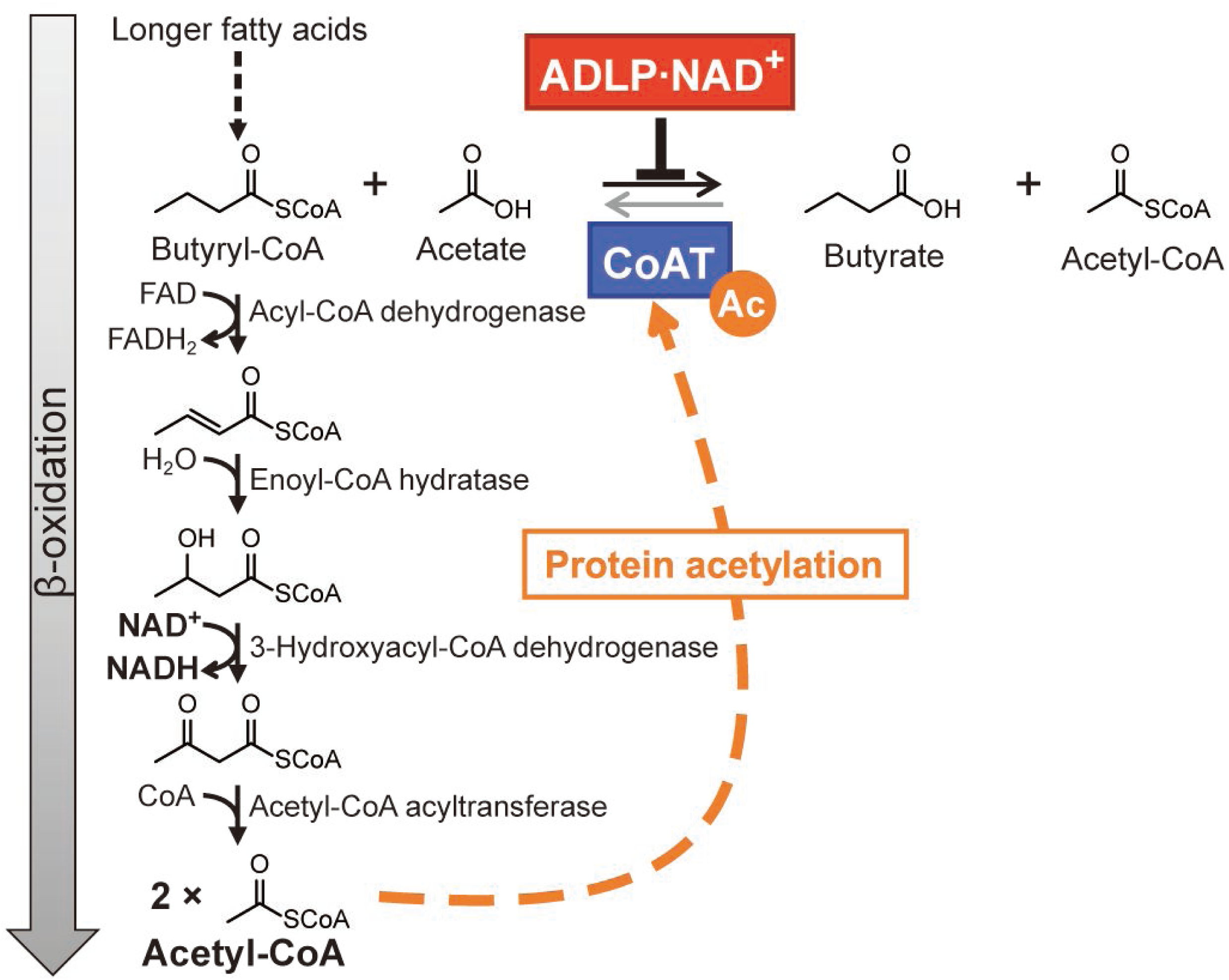
Physiological function of TtCoAT. Reaction schemes of the β-oxidation pathway and TtCoAT from/to butyryl-CoA.

In the present study, we found that CoAT was regulated by both protein acetylation and interactions with catalytically inactive alanine dehydrogenase homolog in *T. thermophilus*. In addition to TtCoAT, *T. thermophilus* possesses several ACS homologs in which the catalytically important lysine residues for acetylation are conserved. Although our previous acetylome study did not detect the acetylation of the conserved lysine residue in each ACS homolog, we postulate that the metabolism of SCFAs and acyl-CoAs, including acetyl-CoA, may be regulated in combination with the possible acetylation status of ACS and the control of TtCoAT activity, which was demonstrated in this study. CobB, which is responsible for the deacetylation of ACS from *S. enterica*, is an NAD^+^-dependent sirtuin-type deacetylase (Starai *et al*., 2002) and a deletion mutant of *cobB* cannot grow on acetate medium because ACS activity cannot be restored due to a defect in deacetylation activity (Starai & Escalante-Semerena, 2004). These studies suggest that not only acetyl-CoA availability, but also the NAD^+^/NADH ratio controls ACS activity in *S. enterica*. Unlike CobB from *S. enterica*, TT_C0104, which is responsible for the deacetylation of CoAT, is a metal-dependent KDAC. In *T. thermophilus*, ADLP, instead of a sirtuin-type KDAC, appears to play a key role in sensing the NAD^+^/NADH ratio in order to maintain metabolic homeostasis through the regulation of CoAT. Importantly, the regulation of CoAT remains largely unexplored, but our findings indicate that CoAT may function as a critical control point in cellular metabolism, responding to key metabolic indicators such as the NAD^+^/NADH ratio and acetyl-CoA availability. Although the distribution of ADLP and the conservation of acetylated lysine residue in TtCoAT appear limited, it is conceivable that in other organisms, CoAT activity may be modulated through alternative mechanisms that integrate NAD^+^/NADH and acetyl-CoA signals to fine-tune metabolic flux. These observations highlight the potential for diverse regulatory strategies within the CoAT family and underscore the importance of CoAT in coordinating cellular energy and acyl-CoA metabolism.

## Methods

### Strains, media, and chemicals

*E. coli* DH5α was used for DNA manipulation, and *E. coli* BL21(DE3) (Novagen, Darmstadt, Germany) and BL21-Codon Plus (DE3)-RIL (Agilent Technologies, CA) were used as hosts for recombinant protein production. 2 × YT medium (Sambrook & Russell, 2001) was generally used for the cultivation of *E. coli* cells and antibiotics and isopropyl-β-thiogalactopyranoside (IPTG) were added to the medium when necessary. TM medium (0.8% tryptone, 0.4% yeast extract, 0.2% NaCl, 1.5 mM MgCl_2_, and 1.5 mM CaCl_2_) and MM medium (Oshima & Imahori, 1974) were mainly used in the cultivation of *T. thermophilus* HB27 and its derivatives. All chemicals were purchased from FUJIFILM Wako Pure Chemicals (Osaka, Japan), Sigma-Aldrich Japan (Tokyo, Japan), or Kanto Chemicals (Tokyo, Japan). We used enzymes from TaKaRa Bio (Shiga, Japan) and TOYOBO (Osaka, Japan) for DNA manipulation.

### Preparation of mutants of T. thermophilus HB27

We selected ten genes (*TT_C0436*, *TT_C0518*, *TT_C0848*, *TT_C0955*, *TT_C1210*, *TT_C1703*, *TT_C1711*, *TT_C1810*, *TT_C1890*, and *TT_C1955*) as candidates for genes encoding protein acetyltransferases by searching the KEGG database using acetyltransf_1 motif (Pfam: 00583) as a query. Each gene was knocked out by replacement with the kanamycin resistance gene, *htk* (Hoseki *et al*, 1999), using pBlueScript KS II (+) (Agilent Technologies, CA) as a vector. The homologous flanking regions (700 bp) of each gene were amplified by PCR using the appropriate primers (primer information is listed in Supplementary Table S2). The *htk* gene was inserted between these flanking regions and ligated into pBlueScript KS II (+). The resulting plasmids were used for the transformation of *T. thermophilus* HB27 (Koyama *et al*, 1986) to generate strains lacking each acetyltransferase gene. Colonies resistant to 50 μg/ml of kanamycin on TM plates were selected, and knockout mutant constructs were confirmed by PCR and Southern blotting. Δ*TT_C1083* (Δ*TtCoAT*) was obtained by replacing *TT_C1083* with the *htk* gene in the same manner.

### Western blotting

Western blotting was performed to detect acetylated proteins. Protein samples were applied to SDS-PAGE and then transferred onto an Immobilon-P PVDF membrane (Merck, NJ) using a Trans-Blot Turbo Blotting apparatus (Bio-Rad Laboratories, Inc., CA). The blot was blocked with 3% (w/v) skim milk in TBS-T buffer (20 mM Tris–HCl, pH 7.6, 150 mM NaCl, and 0.1% Tween 20). To detect protein acetylation, a mixture of rabbit polyclonal anti-acetyllysine primary antibodies (Cell Signaling Technology, MA) and Rockland (Rockland Immunochemicals, PA) was used (1:1,000 dilution) and incubated with the blot at 4°C overnight. The blot was then incubated with goat anti-rabbit IgG-peroxidase antibodies (Sigma-Aldrich, MO) (1:5,000 dilution) at 25°C for 1 h. Signals were detected using a LAS4000 image analyzer (Cytiva, Tokyo, Japan).

### Identification of a highly acetylated protein in T. thermophilus

Protein X, which was identified as a highly acetylated protein in *T. thermophilus*, was purified as follows. *T. thermophilus* was harvested after an overnight culture at 70°C in TM medium. Cells were washed and resuspended with buffer A (20 mM Tris-HCl, pH 8.0, 150 mM NaCl) and then disrupted by sonication. The supernatant prepared by centrifugation at 40,000 × *g* at 4°C for 15 min was fractionated by ammonium sulfate precipitation. Western blotting for precipitation using anti-acetyllysine antibodies showed that 40% saturation ammonium sulfate was sufficient to precipitate Protein X from cell extracts. The precipitant was fractionated using a HiLoad 26/10 Phenyl Sepharose HP (Cytiva) column with 20 mM Tris-HCl pH 8.0 containing 0-1 M ammonium sulfate. The fractions containing Protein X were analyzed by western blotting and applied to a Resource Q (Cytiva) column.

Based on comparisons of SDS-PAGE and western blotting images and samples from the Resource Q column, we estimated the band corresponding to Protein X. Protein X was identified by MALDI-TOF-MS (ultrafleXtreme, Bruker, MA) after in-gel digestion using Trypsin. In-gel digestion was performed as previously reported (Tomita *et al*., 2019). Peak data were searched against the NCBI *T. thermophilus* HB27 database using the MASCOT server (Matrix Science, MA). Protein X with a molecular mass of 50 kDa was identified as TT_C1083 with 79% sequence coverage.

### Preparation of recombinant proteins

The recombinant protein of TT_C1083 (TtCoAT) was prepared with the N-terminal (His)_8_-tag (TtCoATNHis) or C-terminal (His)_6_-tag (TtCoATCHis) by the *E. coli* expression system. The *TT_C1083* gene was amplified with two sets of primers, TT_C1083_Fw_pHis8/TT_C1083_Rv and TT_C1083_Fw/TT_C1083_Rv_Chis6, respectively. Each DNA fragment was then introduced into pHIS8 (Jez *et al*, 2000) using *Bam*HI and *Hin*dIII for the production of TtCoATNHis and into pET26b(+) (Novagen) using *Nde*I and *Eco*RI for the production of TtCoATCHis to yield pHis8-CoAT and pET26-CoATCHis, respectively. The recombinant protein of TT_C1711 was prepared with the N-terminal Strep-tag (TT_C1711NStrep), N-terminal (His)_8_-tag (TT_C1711NHis), and without an affinity tag. The previously constructed plasmid, pHis8-TT_C1711, was used to produce TT_C1711NHis (Yoshida *et al*., 2019). The *TT_C1711* gene was amplified with two sets of primers, TT_C1711_Fw_Strep/TT_C1711_Rv2 and TT_C1711_Fw/TT_C1711_Rv2, respectively. Each amplified DNA fragment was inserted into pBlueScript II KS (+) using *Xba*I and *Eco*RI to confirm their sequences, and was then introduced into pET26b(+) using *Nde*I and *Eco*RI and into pCDFDuet (Novagen) using *Nde*I and *Kpn*I. The resultant plasmids were named pET26-TT_C1711Nstrep and pCDF-TT_C1711, respectively. The recombinant proteins of TT_C1082 (ADLP) and TT_C1770 (AlaDH) were both produced with N-terminal Strep-tag (ADLPNstrep and AlaDHNstrep). Each gene was amplified with two sets of primers, TT_C1082Nstrep_Fw_Strep/TT_C1082Rv and TT_C1770_Fw_Strep/TT_C1770_Rv, respectively. Each DNA fragment was inserted into pACYCDuet-1 (Novagen) using *Nde*I and *Kpn*I to produce plasmids, named pACYCDuet-ADLPNstrep and pACYCDuet-AlaDHNstrep. The nucleotide sequences of the primers used in the present study are shown in Supplementary Table S3. The recombinant proteins of three KDAC homologs (TT_C0104, TT_C1026, and TT_C1954) were prepared as reported in our previous study (Yoshida *et al*., 2019).

*E. coli* BL21-CodonPlus(DE3)-RIL and BL21(DE3) were used as production hosts. *E. coli* cells harboring each plasmid described above were cultured in 2 × YT medium supplemented with appropriate antibiotics (50 μg/ml kanamycin and/or 30 μg/ml chloramphenicol). Gene expression was induced by adding IPTG at a final concentration of 0.1 mM for TtCoATNHis, TT_C1711NHis, and TT_C1711Nstrep and the culture was continued at 25°C for an additional 15 h. To produce TtCoATCHis, ADLPNstrep, and AlaDHNstrep, gene expression was induced by adding IPTG at a final concentration of 1 mM and the culture was continued at 37°C for an additional 15 h. To obtain acetylated TtCoAT (AcTtCoAT) with a C-terminal (His)_6_-tag, *E. coli* BL21(DE3) harboring both pCDF-TT_C1711 and pET26-CoATCHis was used, and gene expression was induced by IPTG at a final concentration of 1 mM at 37°C. Cells were harvested, washed with 20 mM Tris–HCl, pH 8.0, and resuspended in buffer A (20 mM Tris–HCl, pH 8.0, and 150 mM NaCl). Cells were then disrupted by sonication and the debris was removed by centrifugation. The cell lysate was partially purified with a heat treatment at 70°C for 30 min, and the precipitant was removed by centrifugation. The supernatant including the His-tagged recombinant protein was then applied to Ni^2+^-NTA His Bind® Resin (Merck) equilibrated with buffer A and supplemented with 20 mM imidazole, pH 8.0. His-tagged proteins were eluted with buffer A supplemented with 200 mM imidazole, pH 8.0 after washing out contaminated proteins with buffer A supplemented with 20 mM imidazole, pH 8.0. Partially purified proteins, including the Strep-tagged recombinant protein, were applied to Strep-tactin Superflow Plus (Qiagen, Hilden, Germany) resin equilibrated with buffer B containing 50 mM Tris-HCl, pH 8.0 and 150 mM NaCl. The strep-tagged protein was eluted with buffer B supplemented with 2.5 mM desthiobiotin after washing the resin with buffer B. To prepare the TtCoAT·ADLP complex, we used cells producing TtCoATCHis and ADLPNstrep. Proteins partially purified after the heat treatment were mixed and incubated at 70°C for 30 min. Target proteins were purified using Strep-tactin Superflow Plus. Further purification using HiLoad Superdex200 26/600 (Cytiva) with buffer A was conducted if necessary. The purity of each protein was confirmed using SDS-PAGE. Protein concentrations were measured using a protein assay kit (Bio-Rad).

### In vitro (de)acetylation reaction

To examine the TT_C1711-dependent acetylation of TtCoAT, 0.1 mg/ml recombinant TtCoAT was incubated with 0, 20, and 50 μM acetyl-CoA and 50 μg/ml TT_C1711NHis in 50 mM HEPES–NaOH, pH 7.5, in a total volume of 0.1 ml at 50°C for 1 h. After the incubation, the reaction mixture was chilled on ice and mixed with an equal volume of the SDS-PAGE sample buffer to stop the reaction. Each sample (0.5 μg) was applied to 10% SDS-PAGE and acetylation levels were subsequently analyzed by western blotting using anti-acetyllysine antibodies, as described above.

In the enzymatic deacetylation assay, acetylated TtCoAT (AcTtCoAT) was prepared *in vitro* by incubating 0.1 mg/ml recombinant TtCoATCHis with 50 μg/ml TT_C1711Nstrep and 0.2 mM acetyl-CoA in 50 mM HEPES–NaOH, pH 7.5, in a total volume of 10 ml at 50°C for 1 h. AcTtCoAT was purified by the Ni^2+^-NTA column to remove TT_C1711Nstrep and acetyl-CoA. AcTtCoAT was then incubated with each recombinant KDAC homolog (TT_C0104, TT_C1026, or TT_C1954) in the presence or absence of the corresponding co-factors: ZnSO_4_ for TT_C0104 and TT_C1954 and NAD^+^ for TT_C1026. The reaction mixture containing 50 mM HEPES–NaOH, pH 7.5, 0-1 mM ZnSO_4_ or 0-10 mM NAD^+^, 50 μg/ml of each KDAC, and 0.1 mg/ml AcCoAT was incubated at 50°C for up to 2 h. Regarding TT_C0104, reactions in the presence of 5 mM EDTA or 5 mM butyrate were also conducted. The reaction was stopped by immediate cooling on ice and mixing with an equal volume of the SDS-PAGE sample buffer. Samples containing 0.5 μg of TtCoAT were applied to 10% SDS-PAGE and analyzed with western blotting using anti-acetyllysine antibodies.

### Identification of acetylated lysine residues in TtCoAT by nanoLC-MS/MS

To identify acetylated lysine residues, TtCoAT was acetylated by TT_C1711 with acetyl-CoA *in vitro*. The reaction mixture for the acetylation of TtCoAT contained 50 mM HEPES-NaOH, pH 7.5, 200 μM acetyl-CoA, 50 μg/ml of TT_C1711NStrep, and 0.1 mg/ml TtCoATNHis in a total volume of 100 μl. The mixture was incubated at 50°C for 1 h. NH_4_CO_3_ and acetonitrile were added at final concentrations of 50 mM and 10% (v/v), respectively. Proteins were reduced with 20 mM DTT at 56°C for 30 min, and later alkylated with 33 mM iodoacetamide at 37°C for 30 min. Proteins were digested with sequencing grade trypsin (Promega, WI) at a 1:100 enzyme:substrate ratio (w/w) by incubating at 37°C overnight. The final concentration of 0.1% (v/v) trifluoroacetic acid was added and the sample was concentrated by vacuum centrifugation. Trypsin-digested peptides were cleaned with ZipTipC18 (Merck) and then subjected to a nanoLC-MS/MS analysis. MS and MS/MS data were acquired with a Q Exactive mass spectrometer (Thermo Fischer Scientific, MA) as previously described (Kosono *et al*., 2015). MS data were processed using the Proteome Discoverer (ver. 1.4, Thermo Fisher Scientific).

### Site-directed mutagenesis of TtCoAT to identify the main acetylation site

To identify the major acetylation site of TtCoAT, we conducted site-directed mutagenesis using pHis8-CoAT as a template with the corresponding set of primers listed in Supplementary Table S4. TtCoAT derivatives, in which each of the lysine residues at 5, 6, 37, 357, and 401, and both of the lysine residues at 5 and 6 and at 435 and 440 were replaced with arginine, were prepared in the same manner as wild-type TtCoATNHis described above. These TtCoATNHis derivatives (0.1 mg/ml) were acetylated *in vitro* by 50 μg/ml TT_C1711NHis with 50 μM acetyl-CoA. The acetylation levels of each mutant were compared by western blotting using anti-acetyllysine antibodies.

### Activity assay of CoAT

To detect the activity of CoA transferred from acetyl-CoA to SCFAs or carboxylic acid, we analyzed the reaction products by high performance liquid chromatography (HPLC). Fifty microliters of the reaction mixture, which included 100 mM HEPES-NaOH, pH 7.5, 1 mM acetyl-CoA, 10 mM of each carboxylate, and 20 μg/ml TtCoATCHis, was incubated at 50°C for 1 h. After the reaction was stopped by adding 50 μl of 100% methanol, and diluted and pH-adjusted by the addition of 400 μl of 0.5 M MES-NaOH, pH 6.0, 5 μl of the reaction mixture was analyzed using the HPLC system (LC-2000Plus series, JASCO, Tokyo, Japan) equipped with CAPCELL PAK C18 MGII (4.6 × 250 mm, Osaka Soda, Osaka, Japan) at a temperature of 40°C using mobile phase A (water plus 5 mM dibutylammonium acetate) and mobile phase B (100% methanol). Separation conditions were as follows: 10-80% B over 15 min and 80% B for 1 min followed by 10% B for 20 min at a flow rate of 1 ml/min. Enzyme activity was calculated by the decrease in the peak area of acetyl-CoA after the reaction. The difference in the peak area of acetyl-CoA between before and after the reaction using propionate as a substrate was set as 100. We initially attempted to detect activity for propionate, butyrate, succinate, isobutyrate, 3-hydroxybutyrate, valerate, isovalerate, crotonate, hexanoate, heptanoate, acetoacetate, 6-aminohexanoate, pyruvate, α-ketobutyrate, α-ketoglutarate, β-alanine, L-lactate, malate, glutarate, 3-methylglutarate, adipate, anthranilate, 4-aminobenzoate, and palmitate as carboxylates. We then selected substrates exhibiting sufficient activity, which were propionate, butyrate, succinate, valerate, isovalerate, isobutyrate, hexanoate, crotonate, 3-hydroxybutyrate, and heptanoate, for comparisons of activity between TtCoAT and AcTtCoAT.

### Pull-down assay of CoAT

The over-producing strain of *TT_C1083* (Tt27NStrepNHisCoAT) with the Strep-tag and (His)_8_-tag at the N terminus using the constitutive promoter of *slpA* (Fujita *et al*, 2013) was constructed. We constructed pTt-NStrepNHisTT_C1083, which contains the upstream region (700 bp) of *TT_C1083*, the *htk* gene, *slpA* promoter, and 700 bp of a coding region of *TT_C1083* with the Strep-tag and (His)_8_-tag-coding sequences. PCR was performed with appropriate sets of primers listed in Supplementary Table S1 to amplify these DNA fragments. Each of the amplified fragments was ligated into pBlueScript II KS (+) to yield the plasmid pTt-NStrepNHisTT_C1083. The resulting plasmid was used for the transformation of *T. thermophilus* to generate the recombinant *T. thermophilus* strain Tt27NStrepNHisCoAT.

Tt27NStrepNHisCoAT cells were cultured in 800 ml of TM medium supplemented with 40 μg/ml of hygromycin at 70°C for 14 h. Cells were harvested, washed, and then suspended in buffer A for disruption by sonication. The supernatant prepared by centrifugation was loaded onto a column with Strep-tacin resin. The purification of Strep-tagged CoAT was performed in the same manner as that described above. Eluates were concentrated with VIVASPIN 20 (MWCO 10,000, Sartorius, Göttingen, Germany) and subjected to SDS-PAGE. Proteins co-purified with CoAT were identified by in-gel digestion with trypsin followed by a MALDI-TOF-MS analysis as shown above.

### Molecular size analysis of the TtCoAT·ADLP complex

To analyze the oligomeric state or interactions of TtCoAT and ADLP, TtCoATCHis, ADLPNstrep, and the purified TtCoAT·ADLP complex by Strep-tactin resin were subjected to gel filtration chromatography with HiLoad 26/60 Superdex 200 pg equilibrated with buffer A with the flow rate set at 2.5 ml/min. Protein assembly and protein-protein interactions were analyzed by molecular weight calibration using molecular weight markers, gel filtration calibration kits HMW and LMW (Cytiva), and SDS-PAGE of the fractionated eluates.

### Activity assay of alanine dehydrogenase

The activity of alanine dehydrogenase was measured by monitoring the reduction of NAD(P)^+^ to NAD(P)H or oxidation of NAD(P)H to NAD(P)^+^ at 340 nm with a Shimadzu UV2000 spectrophotometer. The reaction mixture for oxidative deamination contained 100 mM potassium phosphate buffer, pH 7.0, 50 mM KCl, 1 mM alanine, and 2 mM NAD^+^ or NADP^+^. The reaction mixture for the reductive amination of pyruvate contained 100 mM potassium phosphate buffer, pH 7.0, 50 mM ammonium chloride, 1 mM pyruvate, and 200 μM NADH or NADPH. One unit of enzyme activity was defined as the amount of enzyme producing 1 μmol of NAD(P)^+^ or NAD(P)H per min.

### Activity assay of TtCoAT·ADLP in the presence of NAD(H) by the DCPIP method

To examine the effects of ADLP and substrates of alanine dehydrogenase on CoAT activity, we used the DCPIP (2,6-dichloropenol indophenol) method in the coupling reaction with CS from *T. thermophilus* HB27 for the detection of acetyl-CoA produced by the CoAT reaction. Recombinant CS was prepared as the fusion with the (His)_6_-tag at the C terminus using *E. coli* harboring the pET26-CSChis vector. We amplified the DNA fragment with the TT_C0978_Fw/TT_C0978_Rv_Chis6 primers using the genomic DNA of *T. thermophilus* HB27 as a template. The amplified DNA fragment was inserted into the pET26b(+) vector to produce pET26b-CSChis. *E. coli* BL21-CodonPlus(DE3)-RIL was used as the expression host. IPTG was added at a final concentration of 0.1 mM to induce gene expression, and the culture was continued at 25°C for 15 h. CS was purified by Ni^2+^-NTA resin in the same manner as that described above.

Acetyl-CoA produced by the CoAT reaction using butyryl-CoA and acetate as substrates was converted into CoA by the CS reaction, and DCPIP was reduced by CoA generated by the reaction. TtCoAT and ADLP, which were purified separately, were mixed at 0.1 mg/ml each and incubated at 70°C for 30 min to prepare the TtCoAT·ADLP complex. The reaction mixture contained 20 mM HEPES-NaOH, pH 7.5, 100 mM acetate, 1 mM oxaloacetate, 0.1 mM butyryl-CoA, 40 μM DCPIP, and/or 1 mM NAD(P)^+^, alanine, or pyruvate. The reaction was started by adding 0.1 μg of TtCoAT or TtCoAT·ADLP and 10 μg of CS, and was monitored by absorbance at 600 nm. One unit of enzyme activity was defined as the amount of enzyme producing 1 μmol of acetyl-CoA. We used various concentrations of butyryl-CoA (0-0.15 mM) for the kinetic analysis. Kinetic parameters were calculated using SigmaPlot12.

### Activity assay of TtCoAT·ADLP in the presence of NAD(H) by HPLC

Since NADH may also reduce DCPIP, it was not possible to confirm the effects of NADH on CoAT activity in the presence of ADLP by the DCPIP method. Therefore, we also analyzed TtCoAT·ADLP activity using HPLC to directly detect the reaction products acetyl-CoA and butyryl-CoA. The reaction mixture with a total volume of 100 μl containing 100 mM HEPES-NaOH, pH 7.5, 1 mM acetyl-CoA or butyryl-CoA, 10 mM butyrate or acetate, 0-1 mM NAD(H)^+^, and 1 μg/ml TtCoAT·ADLP was incubated at 60°C for 15 min. The reaction was stopped by the addition of 10 μl 60% perchloric acid and neutralized by 20 μl of 3 M KHCO_3_. The supernatant (50 μl) prepared by centrifuging was mixed with 200 μl of 0.5 M MES-NaOH, pH 6.0. A small aliquot (5 μl) of the reaction mixture was analyzed using the UHPLC system (Extrema, JASCO) equipped with CAPCELL PAK C18 IF (2.0 × 50 mm, Osaka Soda) with a temperature of 40°C using mobile phase A (water plus 5 mM dibutylammonium acetate) and mobile phase B (100% methanol). Separation conditions were as follows: 10-90% B over 5 min and 90% B for 1 minute at a flow rate of 0.4 ml/min. To calculate the product amount, standard curves were prepared for the peak area of 0-1 mM acetyl-CoA and butyryl-CoA. One unit of enzyme activity was defined as the amount of enzyme producing 1 μmol of acetyl-CoA or butyryl-CoA.

### ITC

To elucidate the affinity of NAD(H) to the TtCoAT·ADLP complex, the calorimetric change was monitored during binding by ITC. This measurement was performed with the iTC200 microcalorimeter (Malvern Panalytical, Malven, UK). We prepared 0.01 mM of the TtCoAT·ADLP complex and 0.5 mM NAD^+^ or 0.25 mM NADH in 20 mM HEPES-NaOH, pH 8.0, and 150 mM NaCl for the cell and syringe, respectively. The concentration of the TtCoAT·ADLP complex was calculated for the decameric complex composed of the TtCoAT tetramer and ADLP hexamer as a unit. NAD^+^ or NADH was injected through the computer-controlled 40-μl microsyringe at an interval of 2 min into the TtCoAT·ADLP solution (cell volume = 200 μl) while stirring at 1,000 rpm. To confirm that NAD(H) bound to ADLP and not to TtCoAT, we measured the calorimetric change during NAD(H) binding to ADLP. In addition to NAD(H), acetyl-CoA, butyryl-CoA, acetate, and butyrate were used as a ligand. Each small compound and ADLP were prepared at 0.2 and 0.02 mM (calculated as a monomeric ADLP), respectively. To quantify *K*_d_ between ADLP and TtCoAT in the presence or absence of NAD(H), we measured the calorimetric change during the titration of 0.1 mM hexameric ADLP to 0.01 mM tetrameric TtCoAT, in the presence or absence of 1 mM NAD(H). All experiments were conducted at 30°C. Enthalpy changes (Δ*H*) and affinities (*K*_a_) upon binding were measured, and the values obtained were used to calculate Gibbs free energy (Δ*G*) and entropy changes (Δ*S*) according to the equations *K*_a_ = e^−Δ*G*/RT^ and Δ*G* = Δ*H* − *T*Δ*S*. Data were analyzed using a single binding site model implemented in the ORIGIN software package provided by Malvern Panalytical.

### Structural determination of CoAT·ADLP by cryo-EM

The recombinant proteins of TtCoATCHis and ADLPNstrep were purified as a complex in the manner described above. Three microliters of the TtCoAT·ADLP complex solution (∼1 mg/mL) was applied to a Quantifoil R1.2/1.3 Cu200 mesh grid (Quantifoil Micro Tools GmbH, Großlöbichau, Germany) that had been glow-discharged for 20 s at 20 mA using a JEC-3000FC sputter coater (JEOL Ltd., Tokyo, Japan). The grid was blotted with a blot force of 0 and blot time of 3 sec in Vitrobot Mark IV (ThermoFisher Scientific, MA, USA) equilibrated at 20°C and 100% humidity and then immediately plunged into liquid ethane. To obtain the structure of the TtCoAT·ADLP complex bound to NAD^+^, NAD^+^ was added to the protein solution at a final concentration of 1 mM. After a 3-min incubation at room temperature, 3 µL of the protein solution was loaded onto the grids and vitrified under the same conditions. The grids were inserted into a CRYO ARM 300 transmission electron microscope (JEOL) equipped with a cold field-emission electron gun operated at 300 kV and an Ω-type energy filter with a 20-eV slit width. Cryo-EM images were recorded with a K3 direct electron detector camera (Gatan, CA, USA) at a nominal magnification of ×60,000, corresponding to an image pixel size of 0.87 Å with a setting defocus range from −0.7 to −2.2 μm, using Serial-EM (Mastronarde, 2005). Holes were detected using YoneoLocr (Yonekura *et al*, 2021). Movie frames were recorded in the CDS counting mode with a total exposure of 3 s and a total dose of ∼80 electrons Å^–2^. Each movie was fractionated into 40 frames. A total of 4,050 and 2,625 movies were recorded for TtCoAT·ADLP with NAD^+^ in two data collections. A total of 3,225 movies were collected for TtCoAT·ADLP without NAD^+^.

Image processing was performed using cryoSPARC (Structura Biotechnology Inc., Toronto, Canada) (Punjani *et al*, 2017). Data sets of TtCoAT·ADLP with and without NAD^+^ were imported, motion-corrected, and used for contrast transfer function (CTF) estimations. A total of 6,056 micrographs from the data set of TtCoAT·ADLP with NAD^+^ and 2,990 micrographs from the data set of TtCoAT·ADLP without NAD^+^ with maximum CTF resolution higher than 5 Å were selected. In the image analysis of TtCoAT·ADLP with NAD^+^, particle picking was performed using a template picker with a particle diameter range of 150 Å. Template images were generated by performing 2D classification in RELION (Zivanov *et al*, 2018) using particle images initially picked with Warp (Tegunov & Cramer, 2019) performed during data collection. In the image analysis of TtCoAT·ADLP with NAD^+^, 2,281,822 particle images were extracted with a box size of 400 pixels (downscaled to 100 pixels via 4× binning). Two rounds of 2D classification into 100 classes were performed, and 1,338,261 particles were selected. Ab initio reconstruction was performed using seven classes, and two of the resulting initial models were selected and subjected to homogeneous refinement. Following homogeneous refinement, 400-pixel box size particle images (downscaled to 360 pixels) were extracted. subjected to homogeneous refinement and subsequent global CTF refinement, local CTF refinement, and non-uniform refinement to give a final map of TtCoAT·ADLP with NAD^+^ with a resolution of 2.26 Å.

In the image analysis of TtCoAT·ADLP without NAD^+^, 1,078,162 particles were selected by the template picker. Template images were generated by the projection of TtCoAT·ADLP with NAD^+^ volume. A total of 630,168 particles were selected by 2D classification and subjected to heterogeneous refinement using three classes. One high-quality class was subjected homogeneous refinement and subsequent global CTF refinement, local CTF refinement, and non-uniform refinement to give a final map of TtCoAT·ADLP in the absence of NAD^+^ with a resolution of 2.69 Å.

To build the atomic model of the TtCoAT·ADLP complex, the crystal structures of TtCoAT and ADLP (see Supplementary Information) were used as the initial models. These structures were fit into the density map as a rigid body and then manually refined using Coot (Emsley & Cowtan, 2004). The model was refined with a function of Real Space Refine in Phenix (Liebschner *et al*, 2019). The iterative cycles of automatic refinement and manual correction were performed to obtain the model with good quality. The model quality was assessed by Molprobity (Williams *et al*, 2018). The statistics of the maps and models were summarized in Supplementary Table S5. Figures were depicted using UCSF Chimera (Pettersen *et al*, 2004) and PyMOL (https://www.pymol.org/).

## Acknowledgments

This work was supported by JSPS KAKENHI (26870113, 17J40083, 20K05804), Amano Enzyme, and Noda Institute for Scientific Research. The cryo-EM analysis was supported by the Research Support Project for Life Science and Drug Discovery (Basis for Supporting Innovative Drug Discovery and Life Science Research (BINDS)) from AMED under Grant Number JP23ama121003. We are grateful to the staff of the Photon Factory for their assistance with X-ray crystallography data collection, which was approved by the Photon Factory Program Advisory Committee (Proposal No. 2015G552, 2017G573, and 2019G546). We thank K. Ohtsuki, and M. Usui in the RRD center of RIKEN CBS for their technical assistance with the nanoLC-MS/MS analysis.

## Data availability

The crystal structures of ADLP bound to NAD^+^ and TtCoAT have been deposited in the Protein Data Bank under the accession numbers 9VAE and 9VAG, respectively. The atomic coordinates and 3D density maps of the TtCoAT·ADLP complex in the NAD^+^-bound and NAD^+^-free forms have been deposited in the Protein Data Bank under the accession numbers 9WNU (NAD^+^-bound) and 9WNS (NAD^+^-free) and in the Electron Microscopy Data Bank under the accession numbers EMD-66124 (NAD^+^-bound) and EMD-66123 (NAD^+^-free), respectively.

